# Multi-omics reveals AKR1B1-regulated galactose metabolic as a driver of gastrointestinal stromal tumor progression

**DOI:** 10.1101/2024.05.21.595125

**Authors:** Xiaonan Yin, Hongxin Yang, Baike Liu, Qinghong Liu, Dan Zhu, Xiaofen Li, Ye Chen, Bo Zhang, Lei Dai, Yuan Yin

**Affiliations:** Gastric Cancer Center, Department of General Surgery, West China Hospital, Sichuan University, Chengdu, Sichuan, 610041, the People’s Republic of China; Department of Gastrointestinal Surgery, The Affiliated Hospital of Guizhou Medical University, Guiyang, Guizhou Province, China; Department of Biotherapy, Cancer Center and State Key Laboratory of Biotherapy, West China Hospital, Sichuan University, Chengdu, 610041, China

**Author notes:** Corresponding author: Yuan Yin, Lei Dai, Bo Zhang.

**Keywords:** GIST, multi-omics, AKR1B1, galactose metabolism, trehalose

## Abstract

The underlying mechanism of malignant progression in gastrointestinal stromal tumors (GISTs) is not fully understood. Despite recent advancements, a comprehensive profile of metabolome, transcriptome, and proteome of GISTs is lacking. This study conducted an integrated multi-omics analysis of GISTs across different risk classifications. By integrating metabolomics, transcriptomics, and proteomics, we identify distinct metabolic patterns and associated biological pathways implicated in the malignant progression of GISTs. Moreover, we identified galactose metabolism and the pivotal rate-limiting enzyme AKR1B1 is dysregulated in GISTs progression. AKR1B1 was upregulated and predicted poor prognosis in GISTs. In addition, AKR1B1 knockdown resulted in trehalose accumulation in GIST cells, thereby inhibiting cell proliferation and mitosis. These findings not only enhance our comprehension of the underlying mechanisms governing GIST progression from a metabolic reprogramming standpoint but also furnish prognostic biomarkers and potential therapeutic targets for GISTs.

## Introduction

Gastrointestinal stromal tumors (GISTs) are the most common mesenchymal tumor of gastrointestinal tract. They can develop anywhere along the gastrointestinal tract, with the stomach and small intestine being the most common locations[1]. The main cause of GISTs is gain-of-function mutations in receptor tyrosine kinase KIT (75-80%) and/or PDGFRA (5-10%)[2]. While targeted therapies like imatinib have significantly improved the treatment of GISTs, acquired resistance to these therapies remains a major challenge[3–4]. The malignant potential of GISTs can vary, ranging from benign behavior in small tumors to highly aggressive tumors. However, KIT and PDGFRA mutations are only the initial events in GIST development[5–6], and there are likely other unknown mechanisms driving the malignancy of the tumor. Current clinical guidelines use various clinicopathological parameters, such as tumor location, size, mitotic rate, and presence of tumor rupture, to assess the malignant potential of GISTs. However, no effective biomarker exists to identify patients with localized GISTs who have a high risk of relapse.

To obtain further insight into the GIST epigenome, Niinuma, T. et al.[7] integrated the epigenomic and transcriptional data of GIST-T1 cells treated with a DNA methyltransferase inhibitor and a histone deacetylase inhibitor, and identified hypermethylation of MEG3 is a frequent event and an indicator of poorer prognosis in GIST patients. Namløs, H. M. et al. [8] integrated the genomic and transcriptomic analyses of untreated, resectable high-risk GISTs and showed increased chromosomal instability (CIN) is an intrinsic feature for tumors that metastasize. A quantitative proteomics analysis of GISTs demonstrated the importance of proteome characterization in promoting the understanding of the etiology of GIST progression[9]. However, an integrated large-scale metabolomic, transcriptomic, and proteomic analysis on GISTs is lacking.

In this study, we conducted a mass spectrometry (MS)-based metabolomics, RNA sequencing, and proteomics on 12 GIST tissue samples with different risk classifications. By integrating multi-omics data, we identified the dysregulation of aldo-keto reductase family 1 member B1 (AKR1B1) in galactose metabolism that may drive tumor progression and serve as prognostic markers or therapeutic targets. These findings provide new insights into the role of metabolic reprogramming in GIST progression and the development of personalized treatment strategies for GIST patients.

## Materials and Methods

### Clinical human sample collection

The study protocol received approval from the Research Ethics Board of West China Hospital, Sichuan University, China [Number: 2019 (1135)]. The research was conducted in accordance with the principles outlined in the Declaration of Helsinki. Written informed consent was obtained from each patient participating in the study. We selected gastrointestinal stromal tumors (GISTs) and matched adjacent normal tissues (ANTs) from 12 patients, which were collected from the Biological Specimen Banks at West China Hospital, Sichuan University, China. These samples were used as a discovery set for comprehensive metabolic, transcriptomic, and proteomic profiling analysis. The detailed clinicopathological characteristics of these 12 patients can be found in Supplemental Table 1. In addition, we collected an independent validation set consisting of 251 GIST tissues and 50 ANTs. All patients included in this study were confirmed a diagnosis of GISTs by two independent pathologists, according to the Chinese consensus guidelines for the diagnosis and management of GISTs. Patients with a history of any cancer treatment prior to sample collection, such as TKI therapy, radiotherapy, or chemotherapy, were excluded from the study. Relevant clinical information including age, gender, tumor size, mitotic count, risk classification, and mutational status, was extracted from the medical records of the patients.

### Cell culture

GIST 882 cell line was obtained from the Shanghai Cancer Institute (Shanghai, China) and cultured in DMEM supplemented with 10% fetal bovine serum (FBS, Gibco) and 1% penicillin/streptomycin (Gibco, USA). GIST-T1 cell line was kindly provided by Dr. Jian Li at Peking University Cancer Hospital and Institute (Beijing, China) and cultured in IMDM supplemented with 20% FBS and 1% penicillin/streptomycin. All cell lines were incubated at 37 ℃ in 5% CO_2_.

### Lentiviral infection

Two AKR1B1-specific shRNAs (shAKR1B1 275: 5′-CACCGCCCATGTGTACCAGAATGAGTTCAAGAGACTCATTCTGGTACACATGGGCTTTTTTG-3′; shAKR1B1 426: 5′-CACCGCTCTTCATCGTCAGCAAGCTTTCAAGAGAAGCTTGCTGACGATGAAGAGCTTTTTTG-3′) were constructed by Gene Pharma Corporation (Shanghai, China) using pGPU6/GFP/Neo vector. A scrambled shRNA (5′-CACCGTTCTCCGAACGTGTCACGTTTCAAGAGAACGTGACACGTTCGGAGAATTTTTTG-3′) was used as a negative control (shNC). The infection was conducted according to manufacturer’s instructions. The efficiency of lentiviral infection was determined by western blotting, and stable strains were obtained through puromycin screening.

### Cell viability assay

Cells from each group were seeded into 96-well plates at a density of 1000 cells per well in 100 μL of DMEM supplemented with 10% FBS. Subsequently, 10 μL of CCK-8 reagent (Dojindo, Kumamoto, Japan) was added to each well, and the plates were incubated for 0, 24, 48, and 72 hours as per the manufacturer’s instructions. The absorbance at 450 nm was measured using a microplate reader (MK3, Thermo Fisher, MA, USA).

### Sample preparation for non-targeted metabolomics

A weight of 30 mg of fresh tissue samples was conducted on ice. Then, 400 μL of pre-chilled methanol: water solution (4:1 v/v; containing 20 μg/mL tridecanoic acid as the internal standard) was added to the samples, followed by homogenization using a high-frequency grinding apparatus for 2 minutes at 60 Hz. The homogenized samples were then subjected to ultrasound treatment in an ice bath for 10 minutes. Subsequently, the mixture was centrifuged at 13000 rpm, 4℃ for 10 minutes. A volume of 200 μL of the supernatant was transferred for vacuum drying and stored at -80℃ until further analysis. The dried samples were reconstituted with 300 μL of methanol: water solution (4:1 v/v; containing 15 μg/mL 17:0 Lyso PC), followed by vortexing for 30 seconds. After treatment with ultrasound for 3 minutes, the samples were allowed to stand at -20℃ for 2 hours. Finally, the metabolite samples were centrifuged at 13000 rpm, 4℃ for 10 minutes, and a volume of 150 μL of the supernatant was collected for LC-MS analysis. A quality control (QC) sample was prepared by combining equal volumes (10 μL) of each individual sample. To minimize system errors during the metabolite extraction process, the metabolite samples were processed in a randomized manner.

### Metabolome analysis based on liquid chromatography mass spectrometry (LC-MS)

Metabolomics analysis of processed GIST samples was conducted using a hybrid quadrupole time-of-flight mass spectrometry (QTOF-MS) system X500R, developed by SCIEX (Framingham, MA), equipped with an electrospray ionization (ESI) turboVTM source operated in the positive ion mode. The data was acquired using SWATH acquisition technology, which is a data-independent acquisition (DIA) mode combined with a dynamic background deduction (DBS) mode. This comprehensive approach allows for the fragmentation and collection of all detectable ions from the sample, enabling both TOF-MS and TOF-MS/MS data acquisition. Qualitative and quantitative analysis of metabolites was performed using Sciex O.S. software V 1.5 (SCIEX) and MetDNA (http://metdna.zhulab.cn/). Statistical analysis was conducted using MetaboAnalyst (https://www.metaboanalyst.ca). Principal Component Analysis (PCA) and Partial Least Squares Discrimination Analysis (PLS-DA) were employed on a normalized peak table to investigate potential differences in metabolite profiles between the two groups. Fold changes and p-values (assessed by the student’s t-test) were computed. The differentially expressed metabolites were subjected to modular analysis using the weighted gene co-expression network analysis (WGCNA) and metabolite pathway enrichment analysis using KEGG analysis[10]. Differential metabolites among the groups were identified based on VIP (variable importance value) >1 and a p-value<.05 according to a student’s t-test. These results were visualized using the Pheatmap package in R (v3.3.2).

### RNA sequencing analysis

Total RNA was extracted using TRIzol (Invitrogen, Carlsbad, CA, USA) in accordance with the manufacturer’s instructions. Subsequently, the quantification and evaluation of the extracted RNA were performed utilizing NanoDrop and Agilent 2100 bioanalyzer (Thermo Fisher Scientific, MA, USA). Total RNA-seq libraries were constructed employing TruSeq Stranded Total RNA with Ribo-Zero Gold (Illumina), following the guidelines provided by the manufacturer. The libraries were subsequently subjected to sequencing on the Illumina Hiseq X Ten Platform. All sequencing procedures were carried out by Basebiotech Co., Ltd (Chengdu, China).

### Quantitative mass spectrometry-based proteome analysis

Approximately 300 μg of protein per sample were utilized for proteome analysis. The proteins were enzymatically digested into peptides of suitable length for detection via mass spectrometry, employing trypsin as the enzyme of choice. The resulting peptides were subjected to desalting using tC18 Cartridges (Sep-Pak, Waters), followed by elution utilizing a solution of 80% acetonitrile (ACN) in 0.1% trifluoroacetic acid (TFA). Subsequently, the eluted peptide fraction was dried using a SpeedVac and reconstituted in 10 microliters (μL) of a solution consisting of 0.1% formic acid in water for further analysis. The peptides were then subjected to nano-ultra-performance liquid chromatography-tandem mass spectrometry (nano-UPLC-MS/MS) employing a Dionex Ultimate 3000 nano UPLC system equipped with an EASY spray column (75 μm x 500 mm, 2 μm particle size, Thermo Scientific). A 70-minute gradient ranging from 8% to 46% acetonitrile was employed during the chromatographic separation. The eluted peptides were subsequently ionized and detected using an Orbitrap Fusion mass spectrometer.

### Western blotting

The cultured GIST cells from all groups were collected, washed twice with 1×PBS, and lysed using RIPA lysis buffer. Equal amounts of total proteins were separated using 10% sodium dodecyl sulfate-polyacrylamide gel electrophoresis (SDS-PAGE) and transferred to polyvinylidene fluoride (PVDF) membranes (Millipore, USA). Subsequently, the membranes were blocked in TBS buffer containing 0.1% Tween 20 and 5% defatted milk at room temperature for 2 hours. The membranes were then incubated overnight at 4 °C with primary antibodies against AKR1B1 (1:2000, A22132, ABclonal), and β-actin (1:5000, 81115-1-RR, proteintech). After that, the membranes were incubated with horseradish peroxidase (HRP)-conjugated secondary antibodies for 1 hour at room temperature. The protein expression was measured using the enhanced chemiluminescence (ECL) substrate kit (Thermo Scientific) and the enhanced chemiluminescence detection system. The protein expression levels were normalized using β-actin.

### Immunofluorescence

The cells were trypsinized and seeded into 6-well glass chamber slides at a density of 1×10^5^ cells per well to grow overnight. The slides were washed with 1×PBS and then fixed for 10 minutes at room temperature using 4% paraformaldehyde. After fixing, the slides were washed twice for 10 minutes each with 1×PBS. The cells were then permeabilized with 0.1% Triton X for 10 minutes followed by two washes with 1×PBS. To block non-specific binding, the cells were incubated in 1×PBS containing 10% goat serum for 15 minutes. Subsequently, the cells were incubated overnight at 4 °C with antibodies specific to phospho-histone H3 (Ser10) at a dilution of 1:100 (Cell Signaling Technology). After the primary antibody incubation, the cells were washed twice for 10 minutes each with 1×PBS and then incubated with a secondary antibody conjugated to Alexa Fluor 568 (dilution 1:100, Invitrogen, USA) for 1 hour at 37°C. Following this, the cells were washed twice for 10 minutes each with 1×PBS and stained with DAPI for 10 minutes at room temperature. All the images were captured using a fluorescence microscope (Olympus).

### Immunohistochemical staining of GISTs microarray

A total of 251 GIST specimens and 50 matched ANTs were processed in a tissue microarray (TMA) arrangement. Representative tumor regions were obtained from formalin-fixed, paraffin-embedded primary cancer tissues. Cores measuring 1.5 mm were extracted from each tumor block and arrayed. Immunohistochemistry was performed on 5 μm sections of the TMA. In brief, staining was carried out using a monoclonal antibody against AKR1B1 (dilution: 1:1000; A22132, ABclonal) in conjunction with a secondary antibody. Areas displaying positive AKR1B1 expression were observed under a microscope, and the number of immunoreactive cells was quantified using Image J software.

### Statistical analysis

All statistical analyses were conducted using SPSS (version 21.0 for Windows, SPSS Inc., Chicago, IL, USA) and GraphPad Prism 9 (GraphPad Prism 9.0, San Diego, CA, USA). Continuous variables are presented as mean ± standard deviation (SD) and were analyzed using the student’s t-test or one-way analysis of variance. Categorical variables are expressed as percentages, and statistical significance was assessed using the Chi-squared or Fisher’s exact tests. Pearson’s or Spearman’s coefficients were calculated to explore correlations between variables. Survival curves were compared using Kaplan-Meier methods and the log-rank test. A p-value < 0.05 was considered statistically significant.

## Results

### Constructing a multi-omics atlas for GISTs

To comprehensively interrogate the metabolic and transcriptional profiles of GISTs, we generated a multi-omics atlas for GISTs with different risk classifications. In total, 12 fresh surgical resections of GISTs and matched ANTs were obtained at West China Hospital, including 6 paired low-risk (LR) GISTs and 6 paired high-risk (HR) GISTs. Upon examining the PCA score plot, it was observed that the replicate samples within each group clustered closely together, indicating a high degree of similarity among the features. However, when comparing between malignant GIST tissues and matched ANTs, distinct characteristics were observed, suggesting substantial differences in both the metabolome and transcriptome profiles (Fig. 1A-B&D-E). Additionally, diverse metabolic and transcriptional profiles were identified between LR and HR GISTs groups (Fig. 1C&F). These findings provide evidence for the reproducible distinction in metabolome and transcriptome profiles among GISTs with varying classifications of risk, revealing intratumor heterogeneity in different risk classifications of GISTs.

**Fig 1.**
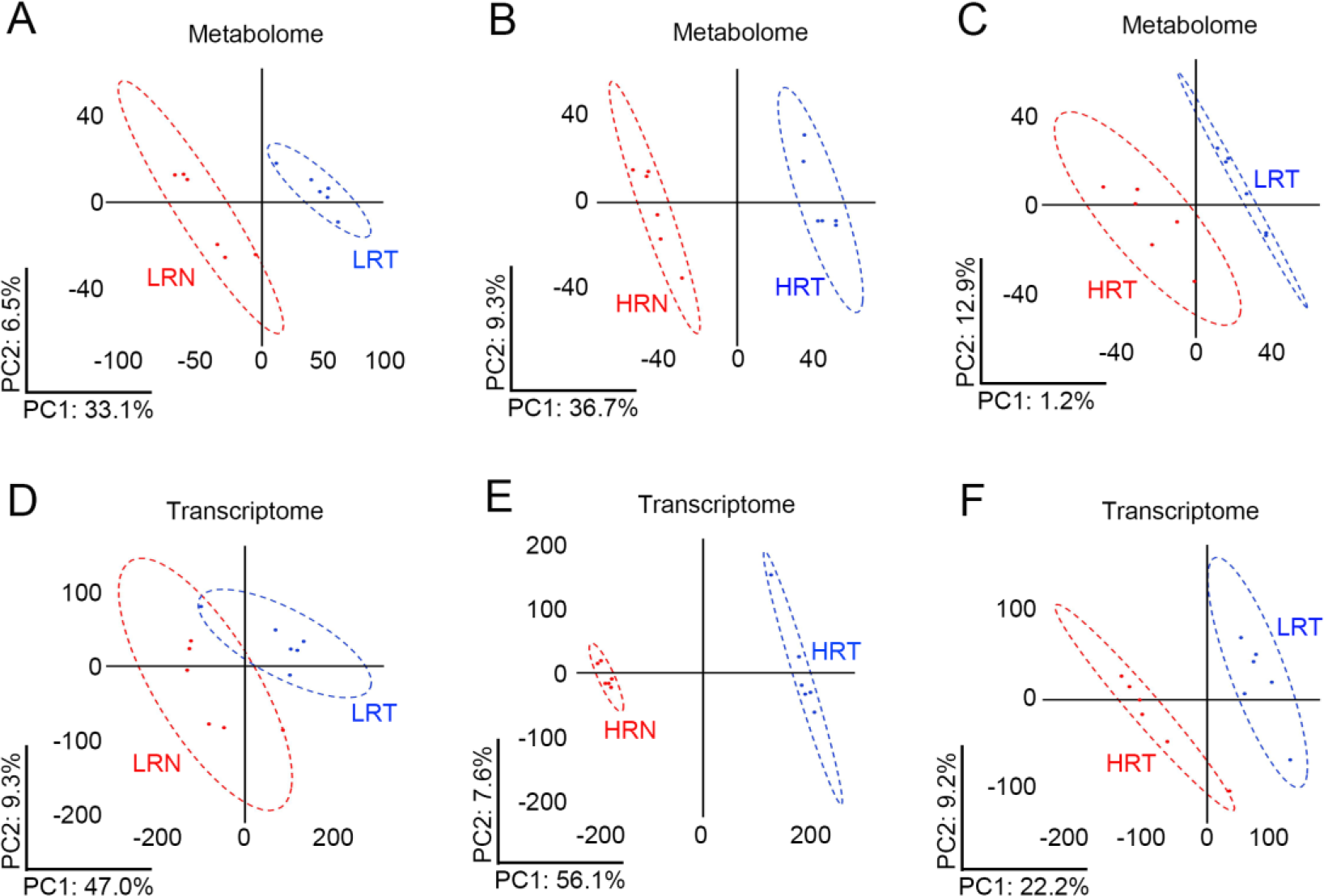
Principal Component Analysis (PCA) of transcriptome and metabolome in GIST with different risk classifications. (A&B) PCA of metabolome in GIST tissues (high- and low-risk) and matched adjacent normal tissues (ANTs); (C) PCA of metabolome in high- and low-risk GISTs; (D&E) PCA of transcriptome in GIST tissues (high- and low-risk) and matched adjacent normal tissues (ANTs); (F) PCA of transcriptome in high- and low-risk GISTs.

### Recognizing differential metabolic modules in GIST tumorigenesis

To explore the metabolites and related biological pathways involved in GIST tumorigenesis, a co-expression network of differentially expressed metabolites was constructed using WGCNA. Following clustering, five modules were identified between LR GISTs and matched ANTs (Figure 2A). Among these five modules, the magenta module exhibited the strongest positive correlation with the LR GISTs group, with a correlation coefficient of 0.78 and a p-value of 0.003. KEGG pathway enrichment analysis revealed that the magenta module predominantly included metabolites that were significantly overrepresented in pyrimidine metabolism (Figure 2B). On the other hand, the turquoise module displayed the highest negative correlation with the LR GISTs group, with a correlation coefficient of 0.81 and a p-value of 0.001. Further KEGG pathway enrichment analysis demonstrated that the metabolites within the turquoise module were enriched in the mTOR signaling pathway (Figure 2C). These findings suggested that the magenta and turquoise modules were most closely related to tumorigenesis in LR GIST groups. Moreover, eight modules were identified between HR GISTs and matched ANTs through the hierarchical clustering tree (Figure 2D). The blue module emerged as the most positively correlated module (correlation = 0.90, p = 7e-05) and showed enrichment in carbon metabolism (Figure 2E). Conversely, the turquoise module was identified as the most negatively correlated module (correlation = -0.96, p = 6e-07) and exhibited enrichment in ether lipid metabolism. These findings indicated that the blue and turquoise modules were most closely correlated with tumorigenesis in HR GIST groups. Taken together, these results suggested that LR and HR GISTs harbored distinct metabolites in process of tumorigenesis, which played important roles in forming their heterogeneity.

**Fig 2.**
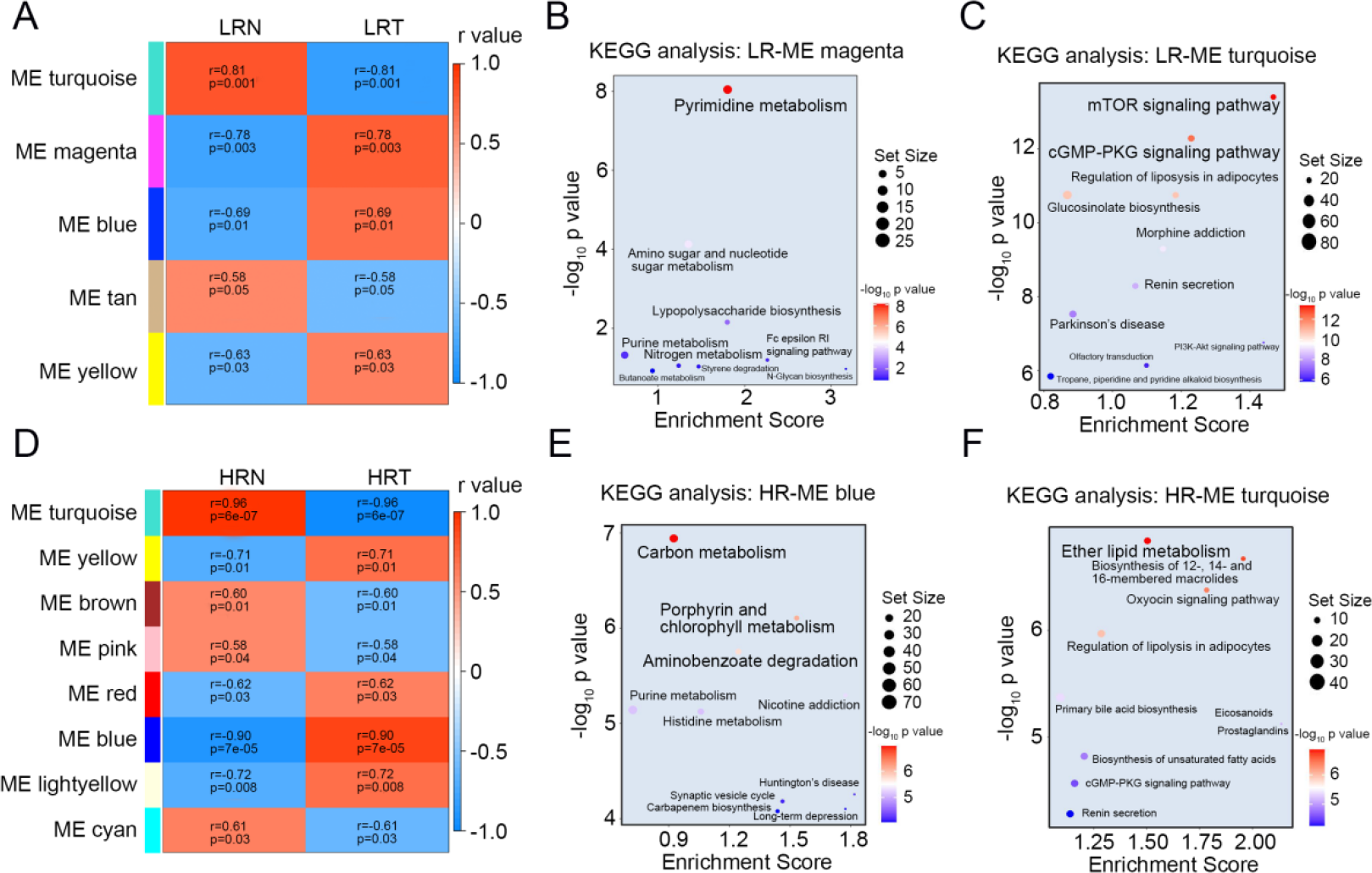
WGCNA of differential metabolic modules in GISTs. (A-C) Differential metabolic modules and KEGG pathways involved in low-risk GISTs and matched ANTs; (D-E) Differential metabolic modules and KEGG pathways involved in high-risk GISTs and matched ANTs.

### Galactose metabolism was identified as malignant-specific metabolic alterations in GIST progression

To identify the key metabolic pathway involved in the malignant progression of GISTs, we conducted Pearson’s correlation analysis to explore the relationship between differentially regulated metabolomes in LR and HR GISTs. The analysis revealed that there was a strong correlation between the LR-ME tan and HR-ME cyan metabolomes, with a p-value of 1e-51 (Fig. 3A). Subsequently, we compared the distribution patterns of LR-ME tan and HR-ME cyan metabolomes between GISTs and their matched ANT tissues using Principal Component Analysis (PCA). The PCA score plot demonstrated that GISTs and ANT tissues could be categorized into two distinct groups (Fig. 3B-C). KEGG pathway enrichment analysis indicated that metabolites present in the LR-ME tan module were enriched in ansamycin biosynthesis (Fig. 3D), while metabolites present in the HR-ME cyan module were enriched in glycerophospholipid metabolism (Fig. 3E). These findings suggest that these pathways may be involved in the overall process of GISTs, including tumorigenesis and malignant progression.

**Fig 3.**
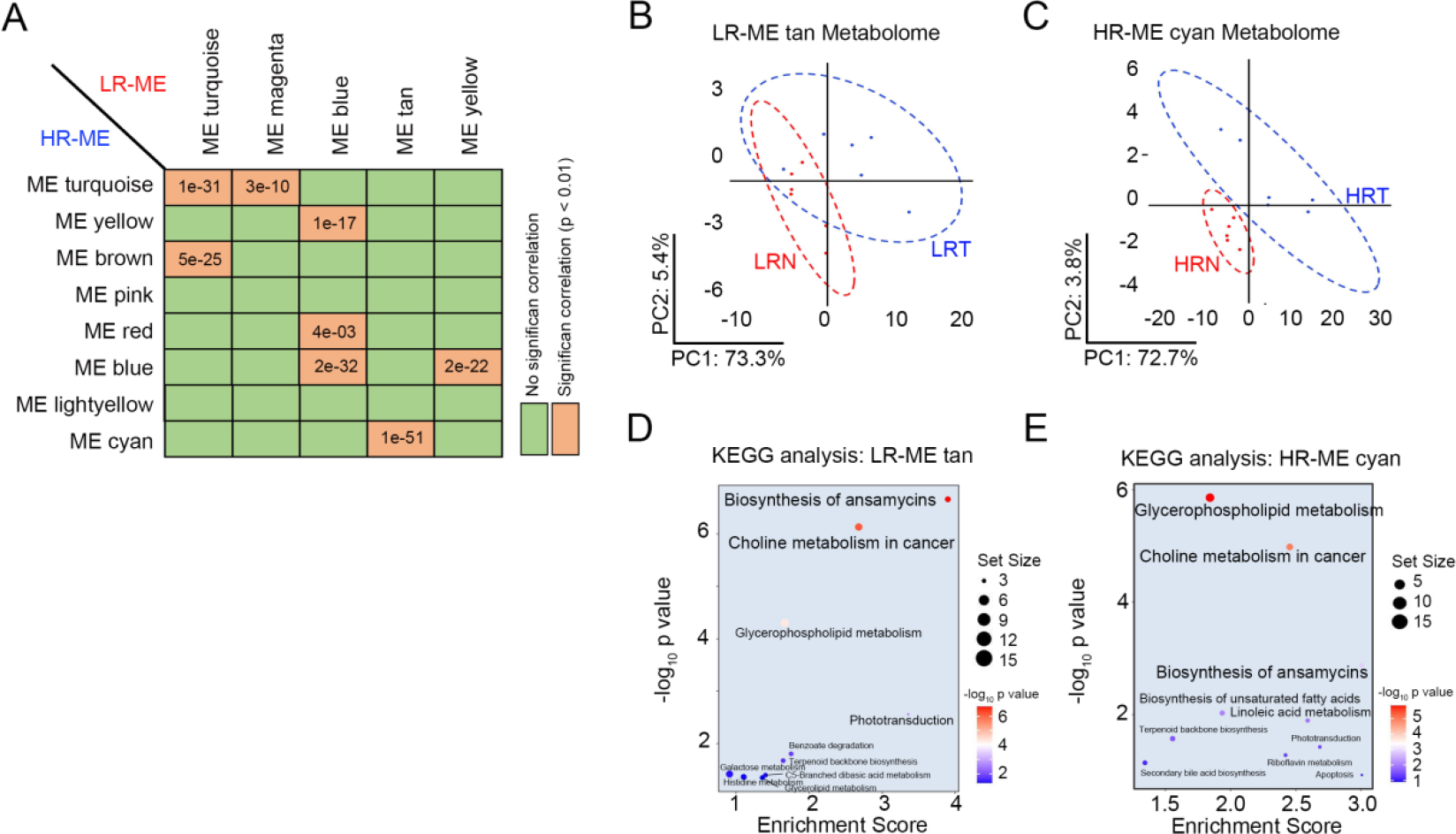
Pearson’s correlation analysis of differential metabolic modules in GISTs with different risk classifications. (A) Pearson’s correlation analysis of differential metabolic modules in high- and low-risk GISTs; (B&D) PCA and KEGG pathway enrichment analysis of LR-ME tan metabolome in low-risk GIST and matched ANTs; (C&E) PCA and KEGG pathway enrichment analysis of HS-ME cyan metabolome in high-risk GIST and matched ANTs.

Next, we identified HR-ME pink and HR-ME lightyellow metabolome were specifically expressed in HR GISTs by Pearson’s correlation analysis (Fig. 3A). Further PCA analysis also revealed distinct metabolic profiles between LR and HR GIST (Fig. 4A&C). KEGG analyses showed that these metabolites, present in HR-ME pink and HR-ME lightyellow modules, were enriched in carbohydrate digestion and absorption, galactose metabolism, as well as phenylalanine, tyrosine, and tryptophan biosynthesis (Fig. 4B&D). These results suggest that these metabolites may play a crucial role in the malignant progression of GISTs. Furthermore, we identified discriminatory metabolites using VIP statistics. Compared to the LR GISTs group, the HR GISTs group exhibited higher levels of key metabolites such as Lacto-N-biose, L-Phosphinothrich, Deoxyinosine, and N-Glucosyinicotinate, while organic acids like N-Succinyl-L-glutamate and 2-Methyl-trans-aconitate were decreased (Fig. 4E&F). Notably, metabolites related to galactose metabolism showed significant changes between LR and HR GIST tissues. These findings suggest that dysregulation of galactose metabolism might contribute to the malignant progression of GISTs.

**Fig 4.**
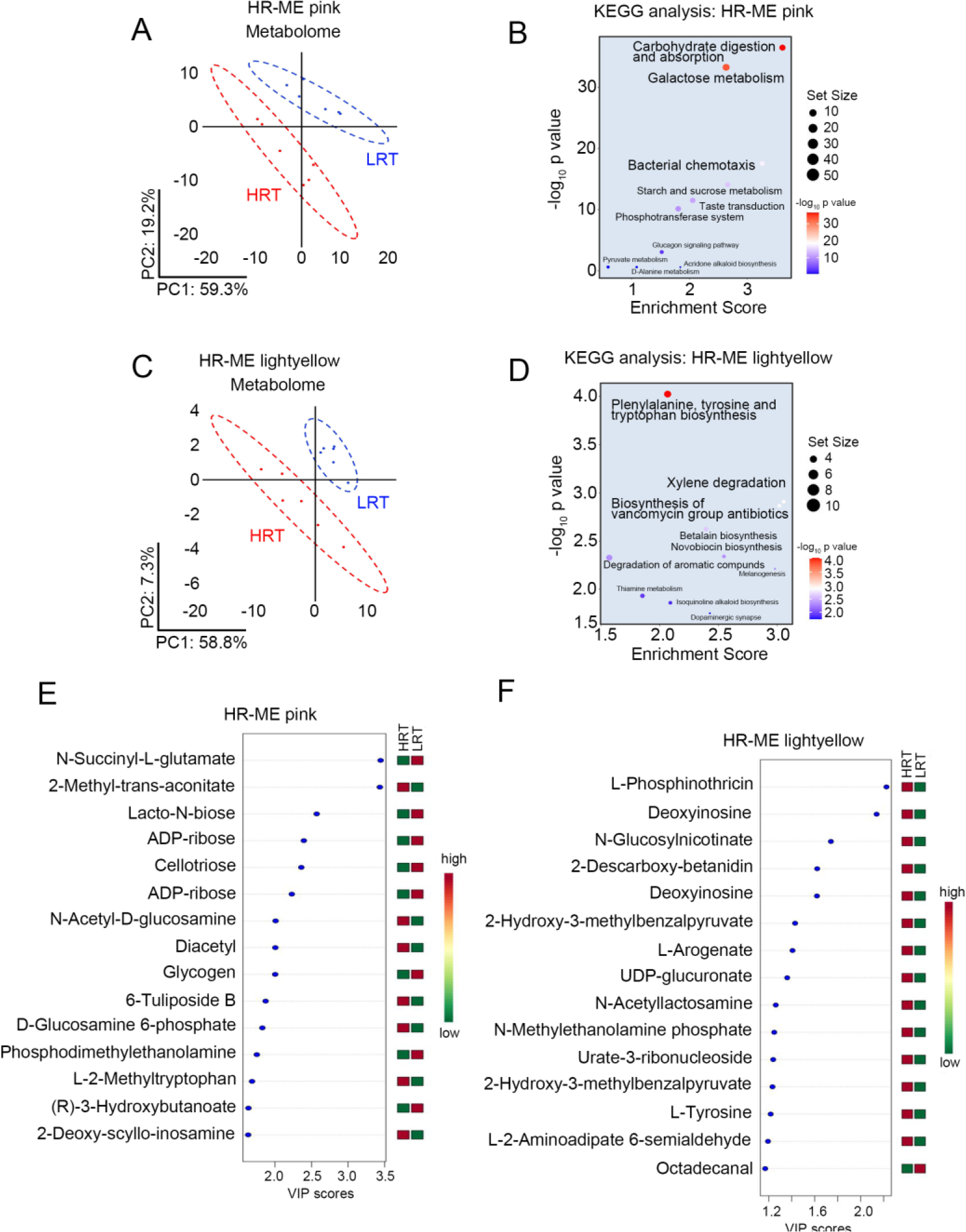
Screening of metabolic modules and KEGG pathways that closely related to malignant progression of GISTs. (A&B) PCA (A) KEGG pathway enrichment analysis (B) and of HR-ME pink metabolome in high-risk GISTs and matched ANTs; (C&D) PCA (C) and KEGG pathway enrichment analysis (D) of HR-ME lightyellow metabolome in high-risk GISTs and matched ANTs; (E) Differential expressed metabolites in the HR-ME pink metabolome determined by the VIP score; (F) Differential expressed metabolites in the HR-ME lightyellow metabolome determined by the VIP score.

### The key rate-limiting enzymes AKR1B1 in galactose metabolism promoted the malignant progression of GISTs

In order to determine the key rate-limiting enzymes involved in galactose metabolism that contribute to the malignant progression of GISTs, a targeted proteomics analysis was conducted. This analysis aimed to quantitatively evaluate the protein levels of Galactokinase-1-Phosphate uridyltransferase (GALT), Galactokinase (GALK1), and aldo-keto reductase family 1 member B1 (AKR1B1) in different risk stratifications of GISTs. The results demonstrated that there were no significant alterations in the expression levels of GALT and GALK1 between LR and HR GIST groups (Figure 5A&B). However, when comparing the LR GIST group to the HR GIST group, a notable increase in the expression level of AKR1B1 was observed in HR GIST group (Figure 5C). These findings suggest that AKR1B1 might involve in the malignant progression of GISTs.

**Fig 5.**
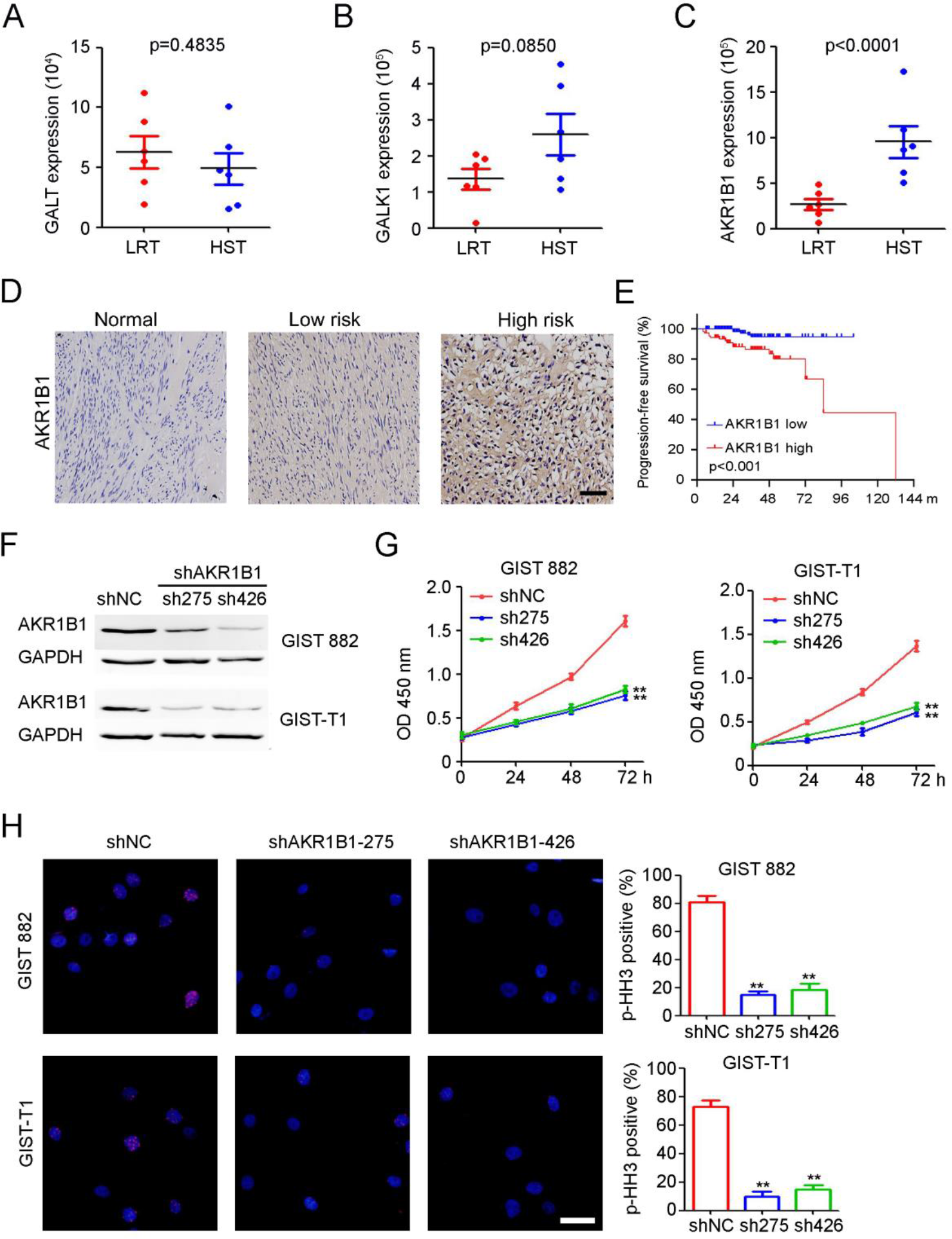
AKR1B1 expression in galactose metabolism was associated with the prognosis of GIST patients and AKR1B1 knockdown inhibited the proliferation and mitosis of GIST cells. (A-C) The expression of GALT (A), GALK (B) and AKR1B1 (C) in low- and high-risk GISTs were detected by targeted proteomics; (D) IHC staining of AKR1B1 expression in tissue microarrays containing 251 GISTs and adjacent normal tissues from GIST patients. Scale bar = 100μm; (E) High AKR1B1 expression was associated with worse progression-free survival in GIST patients; (F) GIST 882 and GIST-T1 cells were infected with lenti-shNC or lenti-shAKR1B1. Stably infected cells were selected by puromycin. Proteins were extracted and subjected to western blotting; (G) Viability of GIST 882 and GIST-T1 cells were detected using CCK-8 assay (n= 5, **p < 0.01); (H) Immunofluorescence analysis of p-histone H3 expression in GIST 882 and GIST-T1 cells, Scale bar = 20μm. Percentage of p-histone H3positive cells in each frame was analyzed (n = 5, **p < 0.01).

To evaluate the expression and potential correlation between AKR1B1 expression and the prognosis of GIST patients, immunohistochemical staining was conducted on tissue microarrays containing 251 GISTs and corresponding adjacent normal tissues. The results exhibited a significant increase in the expression of AKR1B1 in GIST tissues compared to adjacent normal tissues (Figure 5D). Further analysis revealed that AKR1B1 expression was notably higher in HR GISTs compared to LR GISTs (Figure 5D). Based on the median AKR1B1 expression level, the 251 patients were divided into high (112 patients) and low AKR1B1 expression groups (139 patients). Clinicopathological analysis demonstrated a positive association between AKR1B1 expression and tumor size, mitotic count, and the modified NIH risk classification (Table 1). Kaplan-Meier analyses indicated that higher AKR1B1 expression was significantly correlated with worse progression-free survival (PFS) (Figure 5E). Furthermore, to investigate the biological function of AKR1B1 in GIST, shRNA knockdown experiments were carried out using two different shRNAs targeting AKR1B1 in GIST 882 and GIST-T1 cells (Figure 5F). Knockdown of AKR1B1 led to a significant decrease in the proliferation of GIST 882 and GIST-T1 cells compared to the non-targeting shRNA control (NC) (Figure 5G). Additionally, AKR1B1 knockdown resulted in a significant reduction in the mitotic index of GIST 882 and GIST-T1 cells (Figure 5H). Similar findings were observed in GIST cells treated with the AKR1B1 inhibitor, epalrestat (Figure S1). These findings suggested that AKR1B1 promote tumor progression and could potentially serve as a candidate target for tumor therapy in GIST patients.

**Table 1.** Association between AKR1B1 expression and the clinicopathological characteristics of GIST patients.

| Variables | AKR1B1 expression |  | P value |
| --- | --- | --- | --- |
|  | Low expression<br>(n=139) | High expression<br>(n=112) |  |
| Gender (n, %) |  |  |  |
| Male | 51 (36.7) | 54 (48.2) | 0.072 |
| Female | 88 (63.3) | 58 (51.8) |  |
| Age (years), mean ± SD | 57.7±11.7 | 59.0±10.8 | 0.360 |
| Tumor size (cm), mean ± SD | 5.6±2.9 | 7.9±5.3 | <0.001 |
| Mitotic count (n, %) |  |  |  |
| ≤5/50 HPFs | 93 (66.9) | 58 (51.8) | 0.01 |
| 6-10/50HPFs | 28 (20.1) | 23 (20.5) |  |
| >10/50HPFs | 18 (12.9) | 31 (27.7) |  |
| NIH risk classification (n, %) |  |  |  |
| Very low to low | 57 (41.0) | 30 (26.8) | 0.001 |
| Intermediate | 50 (36.0) | 31 (27.7) |  |
| High | 32 (23.0) | 51 (45.5) |  |
| Mutational status (n=130, %) |  |  |  |
| KIT exon 9/11 | 58 (87.9) | 54 (83.1) | 0.397 |
| PDGFRA exon 18 | 3 (4.5) | 7 (10.8) |  |
| WT | 5 (7.6) | 4 (6.2) |  |

### AKR1B1 inhibition resulted in trehalose accumulation in GISTs

To investigate the mechanism underlying AKR1B1’s regulation of GIST cell proliferation and mitosis, a targeted metabolomic approach was employed using MS-based quantification of 32 glucose metabolites in GIST-T1-shNC and GIST-T1-shAKR1B1 cells (Supplemental Table 2). The results revealed significant changes in three metabolites upon AKR1B1 knockdown: increased trehalose, decreased fructose, and D-xylulose (Figure 6A&B). Notably, trehalose emerged as a novel autophagy inducer with reported protective properties in various diseases and tumors. KEGG enrichment analysis indicated that the differentially expressed metabolites were mainly enriched in pathways involving starch and sucrose metabolism, pentose and glucuronate interconversions, metabolic pathways, and ABC transporters (Figure 6C). To assess whether trehalose could inhibit GIST cell proliferation, GIST cells were treated with varying concentrations of trehalose. The CCK8 assay demonstrated a gradual decrease in cell viability with increasing trehalose doses (0 to 10 mM) (Figure 6D). Based on these results, we selected 5 mM trehalose treatment for 24 hours as the condition for subsequent experiments, as the viability of GIST cells treated with 5 mM trehalose did not significantly differ from those treated with 10 mM trehalose. To elucidate the role of trehalose in the mitotic index of GIST cells, p-histone H3 immunofluorescence was performed. The findings revealed a significant decrease in the mitotic index of GIST cells treated with trehalose (Figure 6E). Furthermore, MDC staining was employed to analyze the autophagic process. As illustrated in Figure 6F, the trehalose treatment group exhibited a higher number of MDC-positive staining cells with intense staining intensity compared to the control group. These data collectively indicate that AKR1B1 inhibition resulted trehalose accumulation, which can suppress the malignant progression of GISTs by inhibiting cell proliferation and mitosis, and promoting autophagy of GIST cells.

**Fig 6.**
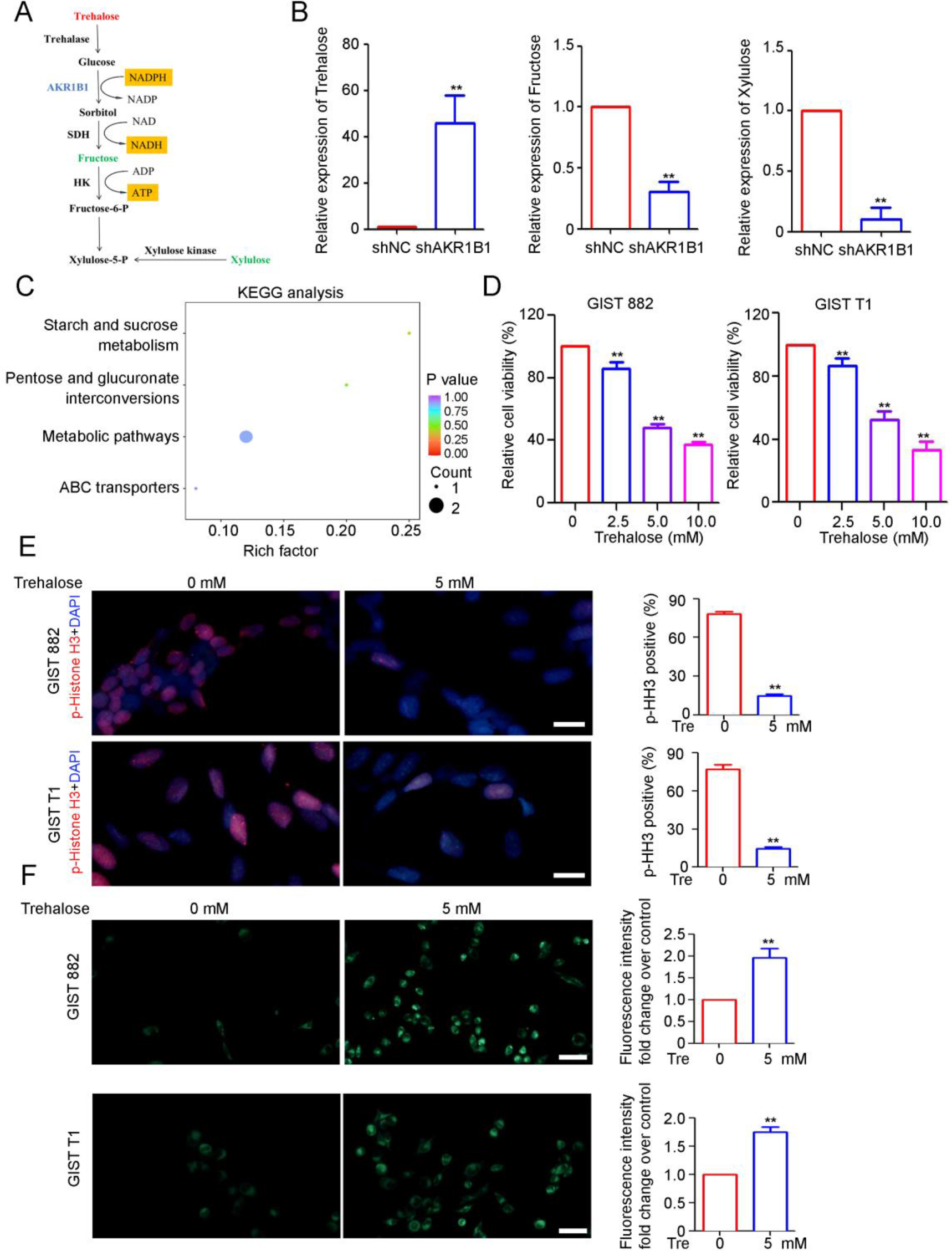
AKR1B1 knockdown inhibited the malignant progression of GISTs by inducing trehalose accumulation. (A) Effect of AKR1B1 inhibition on glucose metabolism in GIST-T1 cells. Metabolites marked in red: abundance increase in resistant versus parental cells, green: decrease, black: unaltered, n = 6 per group; (B) The expression of trehalose, fructose and D-xylulose in GIST cells detected by targeted metabolomics; (C) KEGG pathway enrichment analysis of the differential expressed metabolites after AKR1B1 knockdown; (D) Viability of GIST 882 and GIST-T1 cells after treatment with different doses of trehalose (TRE) detected using CCK-8 assay (n = 3, **p < 0.01); (E) Immunofluorescence analysis of p-histone H3 expression in GIST 882 and GIST-T1 cells after TRE treatment. Percentage of p-histone H3positive cells in each frame was analyzed (n = 5, **p < 0.01); (F) Autophagy in GIST 882 and GIST-T1 cells after TRE treatment was detected by MDC staining (n = 3, **p < 0.01).

## Discussion

The modified NIH risk classification system classify GISTs into very low-, low-, intermediate-, and high-risk categories. The treatment strategies for GISTs range from simple observation to a series of surgical and targeted TKI therapy. GISTs with different risk classifications, despite having similar genetic backgrounds, can display significantly diverse phenotypes. However, a comprehensive multi-omics analysis on GISTs is still lacking.

Metabolic reprogramming is considered a key characteristic of cancer and plays a crucial role in tumor growth, survival, progression, and resistance to therapy. A previous study explored the metabolic landscape of untreated GISTs and identified aspartate as an important “oncometabolite” that is enriched via the TCA cycle and metabolism of alanine, aspartate, and glutamate[11]. Another study by Zhao et al. [12] demonstrated that GISTs with succinate dehydrogenase (SDH) deficiency have elevated levels of succinate, which promotes tumorigenesis. Huang et al.[13] investigated the role of energy metabolism in imatinib-resistant GISTs and found that resistant cells exhibit a higher metabolic activity compared to sensitive cells, while some resistant cells may have reduced oxidative phosphorylation. In this study, we conducted an integrative analysis of metabolic, transcriptional, and proteomic data across different risk classifications of GISTs. Through this analysis, we identified galactose metabolic reprogramming is involved in the malignant progression of GISTs, which could be used to personalize therapy for patients.

Galactose, as a vital nutrient and metabolite, plays essential biological and structural roles in organisms. Recent studies have demonstrated that the reprogramming of galactose metabolism is closely involved in the tumorigenesis and progression of several malignancies[14–16]. For instance, Brown et al.[17] reported significantly decreased expression of sorbitol, a metabolite in the galactose metabolism pathway, in colorectal cancer tissues compared to adjacent normal tissues, implying an inhibitory effect of sorbitol on colorectal cancer tumorigenesis. Shang et al. [18] performed tissue-targeted metabolomic analysis on 50 tissue samples, including 25 papillary thyroid carcinoma (PTC) and 25 healthy thyroid samples, and identified 45 differentially abundant metabolites such as galactinol, melibiose, and melatonin. Their findings confirmed that the galactose metabolism pathway influences PTC development by affecting energy metabolism, with alpha-galactosidase (GLA) being considered a potential therapeutic target for PTC. Furthermore, Han et al.[19] conducted an integrated multi-omics analysis on young breast cancer patients to identify prognostic factors. Their study revealed upregulation of the galactose metabolism pathway in young patients, contributing to worse survival outcomes by regulating cancer stemness. However, the investigation of dysregulated galactose metabolism in GISTs remains limited. Our study, therefore, employed an integrated multi-omics analysis of GISTs across different risk classifications and identified galactose metabolic dysregulation in GISTs, which contributed to the malignant progression of GISTs.

The utilization of galactose requires the collaboration of key rate-limiting enzymes, including galactokinase (GALK), galactose-1-phosphate uridylyl-transferase (GALT), galactose-4-epimerase (GALE), and aldose reductase (AKR1B1). Dysfunction or dysregulation of these enzymes can lead to the accumulation of toxic intermediates in the galactose metabolic pathway, resulting in various diseases and tumors[20–25]. AKR1B1, an important member of the aldo/keto reductase superfamily, is involved in catalyzing the conversion of glucose to sorbitol and has been extensively studied in the context of diabetic pathologies[26–27]. Recent studies have shown that AKR1B1 is overexpressed in several types of tumors, including lung cancer, breast cancer, ovarian cancer, gastric cancer (GC), and colorectal cancer (CRC), and is associated with poor prognosis and drug resistance[25, 28–31]. For instance, Liu et al. [28] reported elevated levels of AKR1B1 in GC tissue, indicating a poor prognosis. They also found that knockdown of AKR1B1 suppressed the AKT-mTOR pathway, leading to the inhibition of proliferation and migration of GC cells. Another study suggested that AKR1B1 plays a role in modulating inflammation and reactive oxygen species (ROS) generation, contributing to the progression from colitis to colon cancer. This highlights the potential of AKR1B1 as a target for the prevention of colitis-associated CRC[31]. However, the role of AKR1B1 in the progression of gastrointestinal stromal tumors (GISTs) has not been investigated to date. In this study, we demonstrated that AKR1B1 was highly expressed in high-risk GISTs and that high AKR1B1 expression was associated with worse prognostic outcomes in GISTs. Knockdown of AKR1B1 resulted in suppressed cell viability and reduced mitotic index in GIST cells. Moreover, the cell proliferation and mitosis can also be inhibited by AKR1B1 inhibitor, epalrestat. These experiments underscore the important role of AKR1B1 in the malignant progression of GISTs and suggest that it could be a potential target for GIST treatment.

Epalrestat, a potent inhibitor of aldose reductase AKR1B1, has been FDA-approved for the treatment of diabetic peripheral neuropathy[32]. Epalrestat has demonstrated significant anticancer effects in breast cancer[33], colorectal cancer[31], and liver cancer[34], both as a standalone treatment and in combination with cytotoxic chemotherapy and targeted therapeutics. Additionally, Kikuya et al. [35] reported that epalrestat enhances the anticancer effects of tyrosine kinase inhibitors (imatinib, nilotinib, or bosutinib) in chronic myelogenous leukemia. This study provides a solid foundation for exploring the application of AKR1B1 inhibitors in combination with imatinib for the treatment of GISTs.

To elucidate the underlying mechanism by which AKR1B1 regulates the proliferation and mitosis of GIST cells, we conducted a targeted metabolomic analysis comparing GIST-NC and GIST-shAKR1B1 cells. We identified a range of differentially expressed metabolites following the knockdown of AKR1B1, including trehalose. Remarkably, trehalose exhibited significant upregulation upon AKR1B1 knockdown in GIST cells. Trehalose, a natural disaccharide derived from glucose, is abundantly found in various organisms such as insects, plants, fungi, and bacteria[36]. It has already been approved for the treatment of several diseases, including ophthalmic and xerophthalmia conditions[37–39]. Notably, recent studies have unveiled trehalose as a potent inhibitor of cancer, exerting its effects by targeting cellular pathways involved in progression, angiogenesis, and metastasis. Molecular-level investigations have revealed its ability to target proteins such as EGFR, PI3K, AKT, VEGF, and MMP 9 within the cell[40]. Our functional analysis of trehalose demonstrated its remarkable capability to suppress cell proliferation and mitosis while inducing autophagy in GIST cells. These findings highlight the potential of trehalose as a therapeutic agent for GISTs.

## Conclusion

To our knowledge, this is the first integrated multi-omics study of GISTs to explore the underlying mechanism of malignant progression in GISTs. Our study identified a series of distinct metabolic patterns and associated biological pathways in the malignant progression of GISTs. We uncovered a dysregulated role for AKR1B1, a rate-limiting enzyme in the galactose metabolic pathway, as a key pathogenic element in GISTs, which was significantly upregulated and correlated with poor prognosis in GISTs. Furthermore, functional studies showed that knockdown of AKR1B1 resulted in trehalose accumulation in GIST cells, thereby inhibiting cell proliferation and mitosis. These results provide a better understanding of the underlying mechanisms accounting for GIST progression from the perspective metabolic reprograming, as well as provide potential therapeutic strategies for the treatment of GISTs.

## Declarations

### Acknowledgements

These authors thank the West China Biobanks, Department of Clinical Research Management, West China Hospital for provision of GIST tissues.

### Author contributions

Xiaonan Yin, Hongxin Yang, Boke Liu, Qinghong Liu and Dan Zhu were involved in acquisition of the data. Yuan Yin and Lei Dai was involved in the study concept and design, and obtained funding. Xiaonan Yin, Bo Zhang and Yuan Yin were involved in the prognosis analysis and patients tissue collection. Xiaofen Li and Ye Chen were involved in analysis and interpretation of the data. Xiaonan Yin, Bo Zhang, Lei Dai and Yuan Yin were involved in drafting of the manuscript and critical revision of the manuscript for important intellectual content.

## Supplementary Information

One supplementary figure and two supplementary tables are included.

---

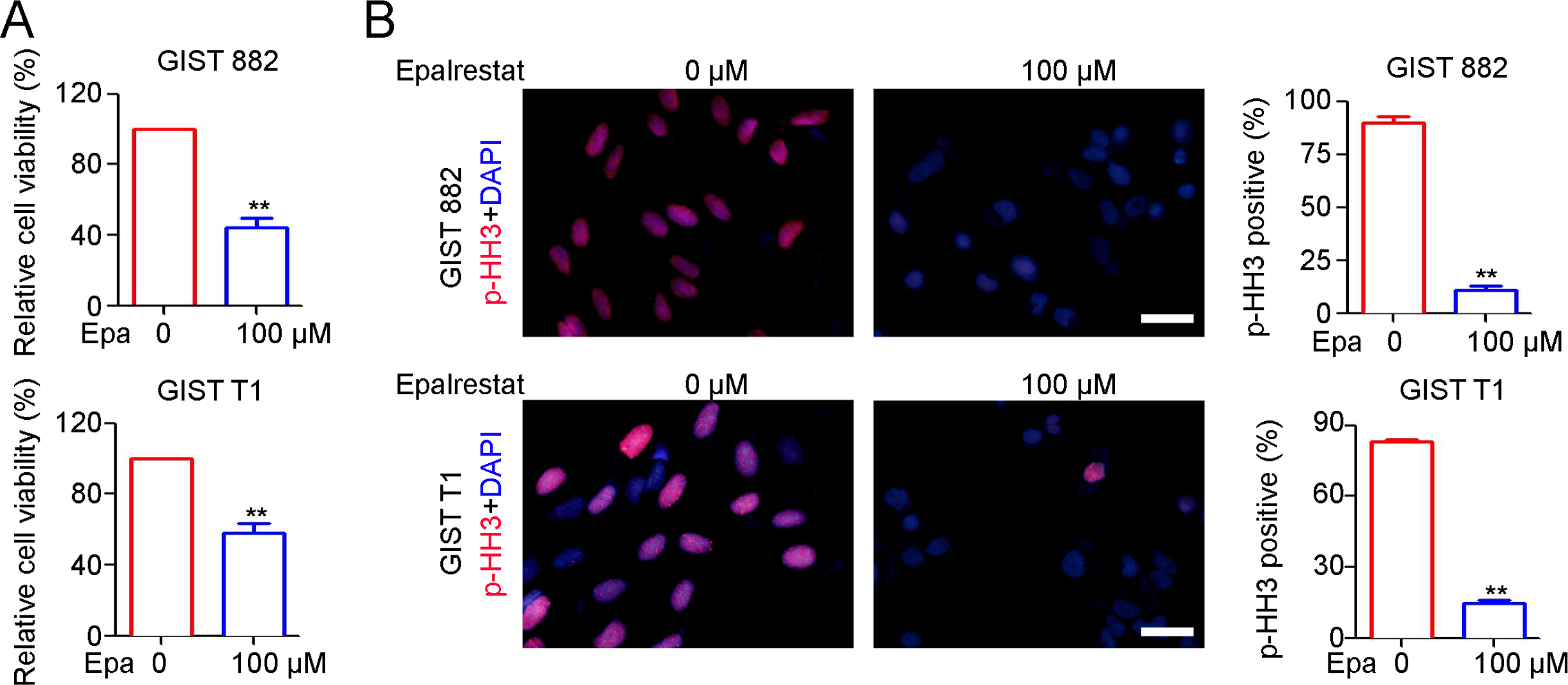

